# Effects of duplicated mapped read PCR artifacts on RNA-seq differential expression analysis based on qRNA-seq

**DOI:** 10.1101/301259

**Authors:** Anna C. Salzberg, Jiafen Hu, Elizabeth J. Conroy, Nancy M. Cladel, Robert M. Brucklacher, Georgina V. Bixler, Yuka Imamura Kawasawa

## Abstract

Best practices to handling duplicated mapped reads in RNA-seq analyses has long been discussed but a gold standard method has yet to be established, as such duplicates could originate from valid biological transcripts or they could be PCR-related artifacts. Here we used the NEXTflex^™^ qRNA-Seq^TM^ (aka Molecular Indexing^™^) technology to identify PCR duplicates via the random attachment of unique molecular labels to each cDNA molecule prior to PCR amplification. We found that up to 64.3% of the single end and 19.3% of the mouse paired end duplicates originated from valid biological transcripts rather than PCR artifacts. For single end reads, either removing or retaining all duplicates resulted in a substantial number of false positives (up to 47.0%) and false negatives (up to 12.1%) in the sets of significantly differentially expressed genes. For paired end reads, only the alignment retaining all duplicates resulted in a substantial number of false positives. This is the first effort to evaluate the performance of qRNA-seq using ‘real-world’ biomedical samples, and we found that PCR duplicate identification provided minor benefits for paired end reads but greatly improved the sensitivity and specificity in the determination of the significantly differentially expressed genes for single end reads.

## Introduction

RNA-sequencing (RNA-seq) has become one of the most popular tools for transcriptome profiling. Most conventional RNA-seq procedures start with random fragmentation of mRNA (or ribosomal RNA-depleted RNA) into short fragments, followed by reverse transcription, and ligation of adapter molecules to the fixed and A-tailed ends. In order to obtain enough material to apply to high-throughput next generation sequencing (NGS), the adapter-ligated cDNA molecules are subjected to 10 –18 cycles of PCR. Under an assumption of linear and non-biased amplification during this PCR step, the users can quantify gene expression, however, it has long been discussed that there is still inevitable skewness in the PCR amplification step that are usually attributed to sequence-dependent bias and underrepresentation of low copy number transcripts. In addition, the conventional RNA-seq usually generates ~20-30% PCR duplicates. The aligner tools such as Tophat or STAR estimates the PCR duplicates when the same sequences are aligned twice, but it is impossible for the aligner to determine whether the duplicates originated from the same or different starting cDNA molecules. Molecular Index^™^ has been conceptualized and developed in gene expression microarray [1], RNA-protein interactions [2] and most recently it was successfully applied to NGS [3,4,5,6,7]. The key concept of Molecular Index^™^ technologies is to add two sets of 96 distinct Molecular Index Adaptors to tag each cDNA molecule with a total of 9,216 (96 × 96) possible combinations of 5′ and 3′ adapters. This allows any two identical molecules to be distinguishable with odds of 9,216/1, and thus the PCR duplicates can be readily distinguished from true biological duplicates. The method has been integrated into a commercially available kit, namely NEXTflex^™^ Rapid Directional qRNA-Seq Kit^™^ [7]. This kit (called “qRNA-seq kit” hereinafter) was elaborated to generate directional RNA-seq libraries, on top of the Molecular Indexing feature, which allows proper detection of antisense and non-coding RNA expression that were largely unavailable due to the overlapping nature of such genes that are transcribed from the different strand but yet from the same/overlapping regions. The qRNA-seq technology itself was developed in 2011, however, to our knowledge, no paper has been published in evaluating the impact of qRNA-seq technology using ‘real-world’ biomedical samples. We aimed to be the first to evaluate the performance of qRNA-seq using ‘real-world’ biomedical samples and propose best practices for handling duplicated mapped reads in RNA-seq analyses, that have long been discussed but a gold standard method has yet to be established.

## Results

### Study design

We used the qRNA-seq technology to identify duplicated mapped read PCR artifacts in 21 mouse and 15 rabbit single end, and 2 mouse paired end samples, with each mouse sample belonging to one of seven phenotypical groups. We first trimmed the qRNA-seq labels and aligned the samples to the mouse or rabbit genome using tophat, a splice junction mapper for RNA-seq reads. The duplicates were then annotated using Broad Institute’s Picard MarkDuplicates. Duplicate reads are defined to be reads that have the same alignment. Specifically, single end reads are considered to be duplicates when they have identical chromosome, start position, sequence length and cigar string, while paired end reads require those fields to be identical in both reads of the template. After alignment, the qRNA-seq labels were recovered, resulting in one label for single end reads, and two labels for paired end reads, from the right and left sequences of the template. PCR duplicates were identified to be those duplicates that had the same qRNA-seq label(s). In addition, to calculate the impact of the second qRNA-seq label in paired end reads, an alternate set of PCR duplicates was extracted in which only the first qRNA-seq label was required to match. For each sample, the RNA transcripts were then assembled and their abundances were calculated using Cufflinks for each of 3 alignments: a) all duplicates were removed, b) all duplicates were retained and c) only PCR duplicates were removed. In each case the mouse and rabbit transcripts were separately merged using Cuffmerge, and mouse groups were compared using Cuffdiff to calculate the significantly differentially expressed genes at thresholds of q < 0.05, q < 0.01 and q < 0.005. We extracted the false positives and false negatives from the sets of significantly differentially expressed genes for both the alignments retaining or removing all duplicates, when considering as the gold standard the corresponding gene set resulting from the alignment having only the PCR duplicates removed. We further characterized the transcripts in the false positive and false negative sets based on transcript length, number of exons and percentage of GC content.

For the mouse study, 7 groups of samples (N=3/group) were harvested from either mouse papillomavirus-infected or non-infected control HSD:NU mice. Group G1 (samples NC15, 16, 17) consisted of non-infected cutaneous tissues, G2 (NC01, 04, 06) of cutaneous lesions, G3 (NC02, 03, 05) of adjacent tissues of the cutaneous lesions, G4 (NC27, 28, 29) of infected tongue, G5 (NC18, 19, 20) of control tongue, G6 (NC24, 25, 26) of infected vaginal samples and G7 (NC21, 22, 23) of non-infected control vaginal tissues. For the rabbit study (samples NC07-14, NC30-36), five lesions were harvested from wild type CRPV, two lesions from four different mutants and two normal tissues at week nine or week 16 post infection. All samples NC01-36 were sequenced in single end mode and samples NC02,06 were sequenced in paired end mode as well.

### Evaluation of consistency in Molecular Indexing

We first examined consistency of qRNA-seq labels (Molecular Index) across all 96 to 96x96 labels and among sequenced samples. On average, 3.19% (0.87-4.56%) of the mouse single end reads and 3.09% (1.32-6.70%) of the rabbit reads did not have a valid qRNA-seq label, and 2.14% of the paired end reads had at least one invalid label. Of the paired end read templates having at least one invalid label, the first read in the template had on average less (39.8%) invalid qRNA-seq labels than the second. Sequences with invalid labels were excluded from all downstream analyses.

The labeling probability of the qRNA-seq labels was calculated by calculating the counts of each qRNA-seq labels by being divided by the average of all counts (“normalized counts” in Figure 1). 96 qRNA-seq labels in the raw sequences of the single end read samples was calculated. For paired end sequences, the distribution of the Read 1 and Read 2 in the templates were calculated separately (Figure 1C) as well as that of the combined 96x96 qRNA-seq labels (Figure 1D). When normalized based on a sample’s average number of valid labels, the distribution of labels for a sample had on average a range of 0.273-1.912 and a standard deviation of 0.333 for the rabbit samples, and a range of 0.246-1.854 and 0.339 standard deviation for the mouse samples, indicating that there was a bias in the qRNA-seq label selection. In fact, the frequency of each of the 96 qRNA-seq labels across the samples had an average standard deviation of only 0.0409 for the rabbit samples and 0.0303 for the mouse samples, indicating that preferential qRNA-seq label selection was uniform across samples. The most frequent qRNA-seq label in the single end sequences was “GGCGTATT,” with 1.91 and 1.85-fold enrichment in the rabbit and mouse samples respectively, and the least frequent one was “TAGCTAGC,” with 3.66 and 4.07-fold depletion. In the paired end sequences, the label combination “ATCGAACC-ATCGAACC” was the most frequent with 3.20-fold enrichment, and “TAGCTAGC-TATAGCGC” was the least frequent with 6.67-fold depletion.

**Figure 1.**
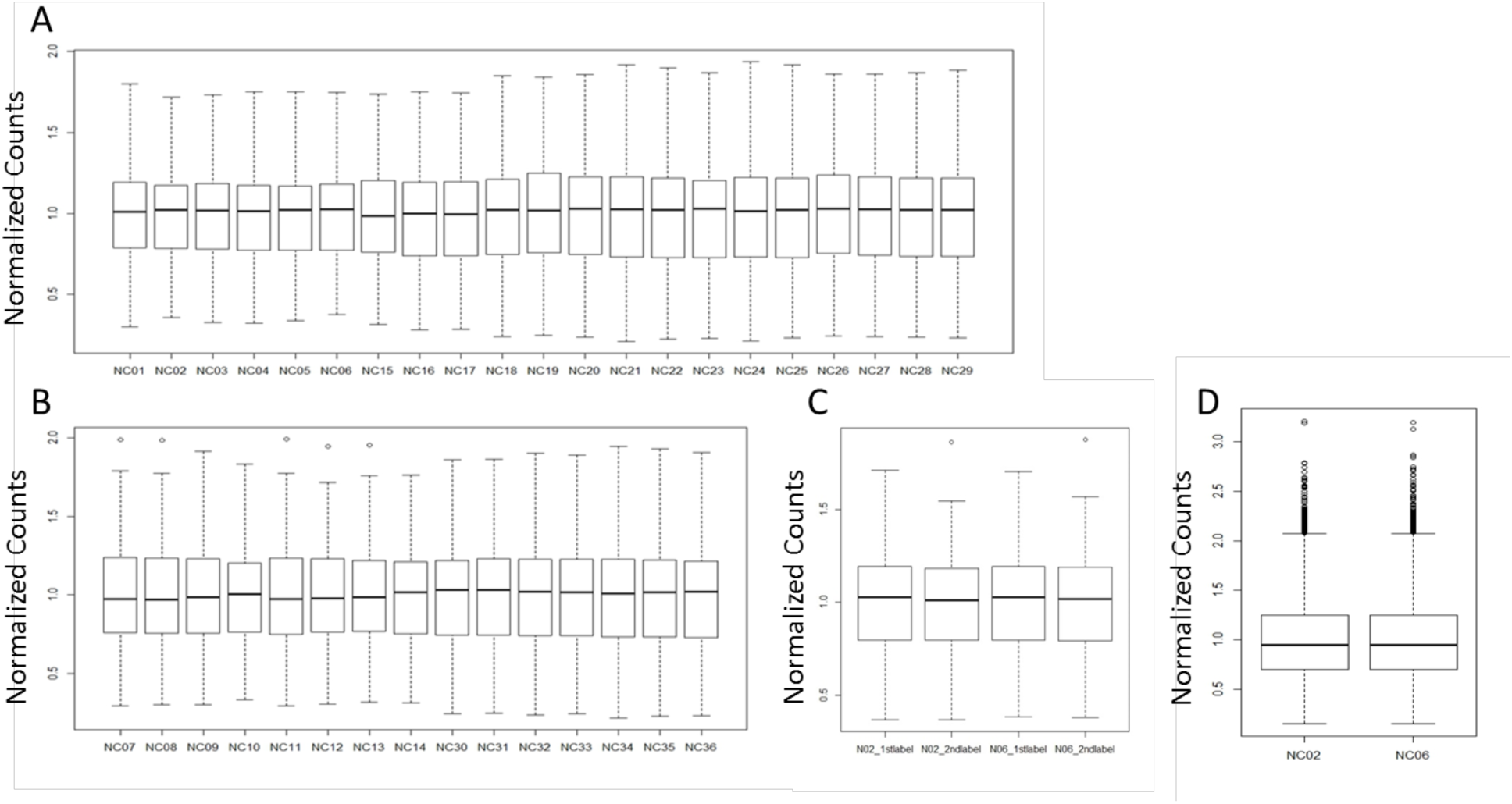
Boxplots displaying the distribution of the normalized qRNA-seq labels in the raw sequences of the (**A**) single end read mouse samples, (**B**) single end rabbit samples, (**C**) first and second reads in the paired end templates calculated separately and (**D**) combined 96x96 qRNA-seq labels in the paired end templates, respectively. The horizontal lines within the box indicate the median, boundaries of the box indicate the 25^th^- and 75^th^ -percentile, and the whiskers indicate the 9^th^- and 91^th^ percentile range of the ratio of each distinct qRNA-seq labels. The labels falling outside the 9^th^- and 91^th^ percentile range are plotted as outliers of the data.

### Amount of non-PCR artifacts (biological duplicates) and minimum advantage by addition of second set of labels in Read 2

The reads were aligned to the appropriate genome, mm10 for mouse or oryCun2 for rabbit samples, with the 8 nucleotide qRNA-seq labels removed, and the alignment duplicates were annotated using Picard MarkDuplicates. For single end reads, an average of 21.2 and 28.0 million reads from the rabbit and mouse samples were mapped to the genome, respectively. For paired end reads, only those alignments with templates having multiple segments in sequencing and where each segment was properly aligned were retained. On average, 22.4 out of 125 million alignments (17.9%) satisfied this requirement and were retained for further analysis. For single end reads the 8 nucleotide qRNA-seq labels of the aligned reads were then recovered, and for paired end reads both labels, from the right and left sequences of the template, were recovered. For single end reads, PCR duplicates were considered to be those alignments marked as duplicates that had identical chromosome, start, end (start plus read length), cigar string and qRNA-seq label to some other aligned read. For paired end alignments, PCR duplicates were defined with an additional requirement that those metrics to be identical in both segments of the cDNA fragment where Read 1 and 2 are aligning to with the same cigar strings and that the fragment to have the same length. On average 37.4% (16.8-59.7%, Figure 2A) of the mouse and 49.7% (10.3-68.9%, Figure 2B) of the rabbit single end duplicates, and 19.3% (Figure 2C) of the paired end duplicates were non-PCR artifacts (i.e. biological duplicates). In addition, to calculate the impact of the second qRNA-seq label in paired end reads, an alternate set of PCR duplicates was calculated in the same way except that the Read 2 qRNA-seq label was not required to match. On average, 15.7% of the paired duplicates were non-PCR artifacts using this alternative PCR definition (Figure 2C). Therefore, the 96x96 labels identified an additional 3.6% of duplicates to be non-PCR artifacts, showing that the much smaller percentage of PCR duplicates in paired end versus single end reads was due mostly to the more stringent definition of read duplication which requires both reads in the template to have identical alignments, rather than the availability of 2 orders of magnitude more label combinations.

**Figure 2.**
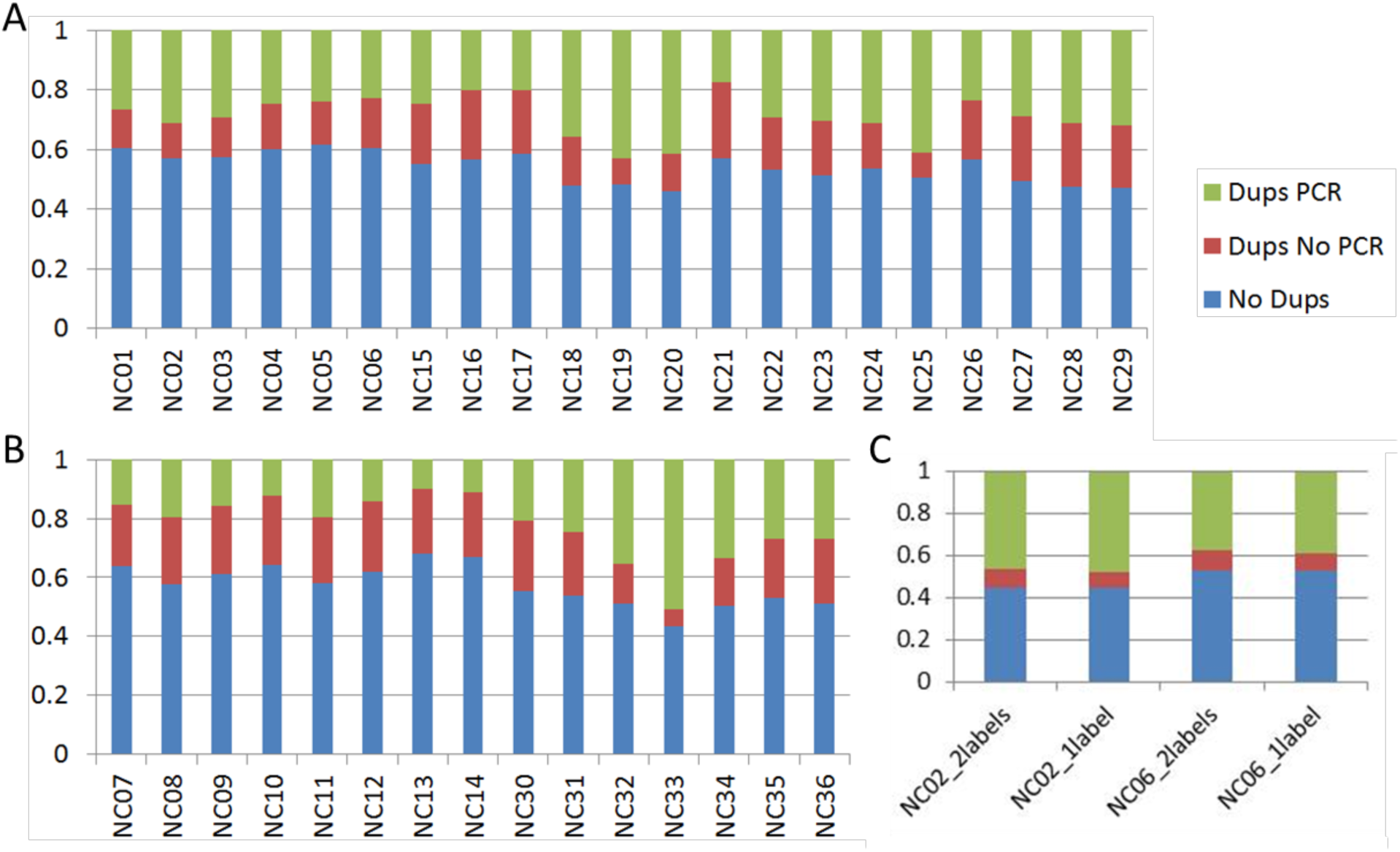
Fraction of PCR duplicates are shown in stacked bar plots. (**A**) and (**B**) represent results from single end reads for (**A**) mouse and (**B**) rabbit projects, respectively. PCR duplicates (“Dups PCR” shown in green bars) were considered to be those alignments marked as duplicates by MarkDuplicates that had identical chromosome, start, end (start plus read length), cigar string and qRNA-seq label to some other aligned read. Averages of 37.4% (16.8-59.7%, **A**) of the mouse and 49.7% (10.3-68.9%, **B**) of the rabbit were non-PCR artifacts (i.e. biological duplicates, “Dups No PCR” shown in red bars) among all duplicated reads (“Dups PCR” + “Dups No PCR”), respectively. (**C**) represents a result from paired end reads of the two identical mouse samples that were analyzed by single end reads as well (NC02 and NC06, **A**). PCR duplicates were defined to have an additional requirement that these MarkDuplicates metrics to be identical in both segments of the template and that the template have the same length (NC02_2labels and NC06_2labels). On average 19.3% were non-PCR artifacts. An alternate set of PCR duplicates was calculated in the same way except that the second qRNA-seq label was not required to match (NC02_1label and NC06_1label). Uniquely aligned reads (“No Dups”) are shown in blue bars.

### Removing or retaining all duplicates contributes to a substantial number of false findings

Each of the 21 mouse single end samples belonged to one of seven phenotypical groups, with each group containing 3 replicates. The 2 mouse paired end samples belonged each to different of such groups. To determine the impact of these findings on downstream analyses, we calculated the significantly (at q < 0.05, q < 0.01 and q< 0.005) differentially expressed genes of each pairwise group comparison for 3 sets of alignments: a) all duplicates retained (Dups, blue+red+green bars in Figure 2 corresponds to pink in Figure 3), b) all duplicates removed (No Dups, blue bars in Figure 2 corresponds to gray in Figure 3), and c) only PCR duplicates removed (No PCR Dups, blue+red bards in Figure 2 corresponds to gold in Figure 3). First, the RNA transcripts for each of such 3 alignments were assembled and their abundances were calculated for each sample. Then the 21 possible pairwise comparisons between the 7 phenotypical groups were calculated separately for each of the 3 sets of alignments and the significantly differentially expressed genes at thresholds of q < 0.05, q < 0.01 and q < 0.005 were extracted for each comparison.

**Figure 3.**
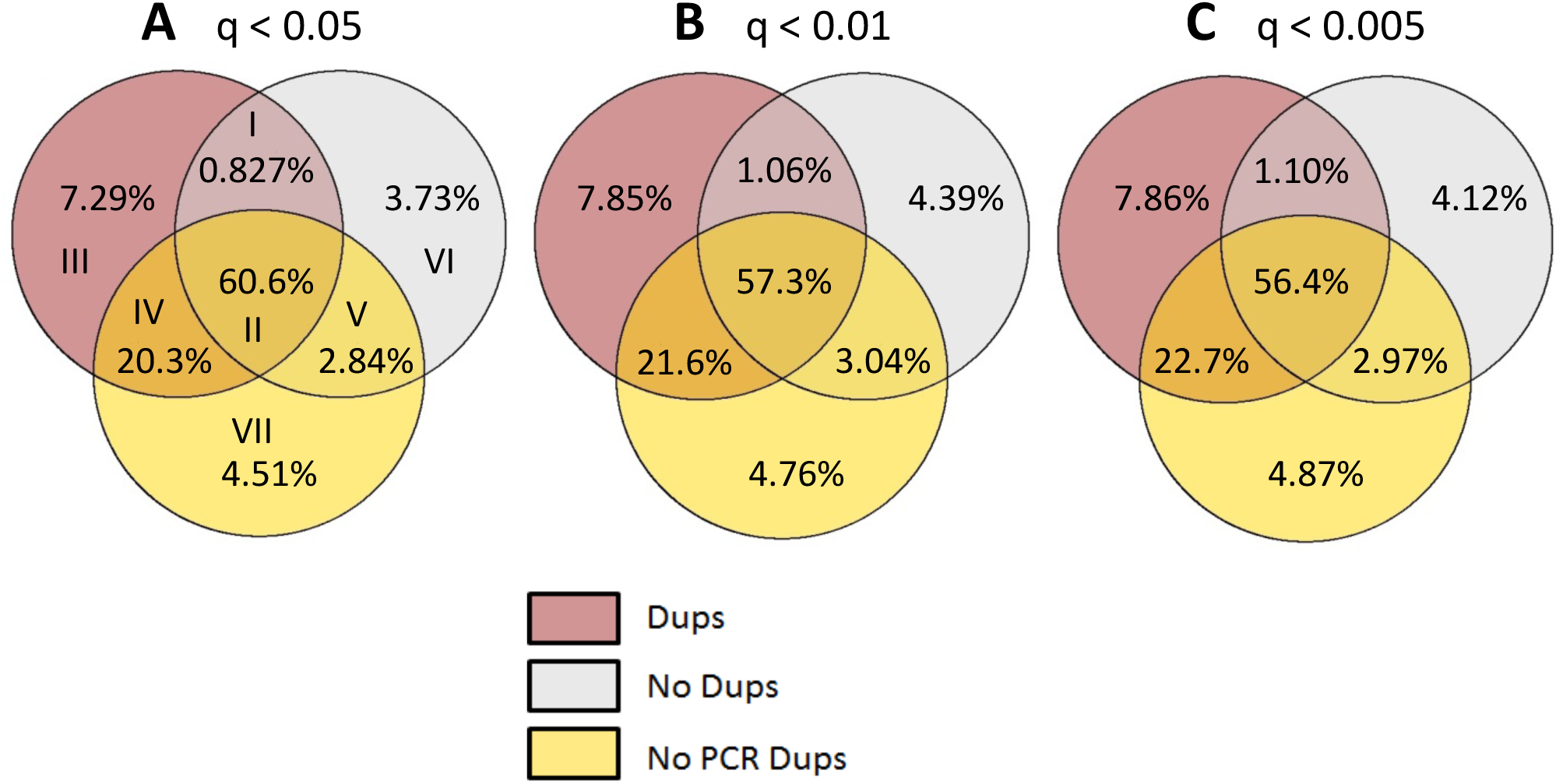
Venn diagrams to show *%* of false positives and false negatives in significantly differentially expressed genes at altering significance threshold, (**A**) q < 0.05, (**B**) q < 0.01 and (**C**) q< 0.005. Average % of significantly differentially expressed genes from each of the 21 pairwise group comparisons was calculated for 3 alignments: all duplicates retained (Dups), all duplicates removed (No Dups) and only PCR duplicates removed (No PCR Dups). Dups false positives are calculated as sum of regions I+III, No Dups false positives are as sum of regions I+VI, Dups false negatives are as sum of regions V+VII, and No Dups false negatives are as sum of regions IV+VII, respectively. All of the 21 pairwise data are available from Supplementary Tables.

For single end reads at a significance threshold of q < 0.05, on average 8.12% (4.60-12.1%) of Dups and 4.56% (2.76-6.80%) of No Dups were false positives and 7.35% (3.86-22.0%) of Dups and 24.8% (15.1-47.0%) of No Dups and were false negatives and when taking the No PCR Dups alignments as the gold standard (Figure 3A). Furthermore, the percentage of false positives and false negatives increased as the significance thresholds were made progressively more stringent (Figures 3B and 3C). All of the 21 pairwise data are available from Supplementary Tables.

For paired end reads, 55% of Dups and 0% of No Dups were false positives and 2.6% of Dups and 2.6% of No Dups were false negatives and at q < 0.05 (Figure 4A), which indicates No Dups approach can be considered as the most reliable method while not using Molecular Indexing, if one can afford paired end analysis. The percentage of false positives, however, increased in No Dups as the significance thresholds were made progressively more stringent (0% vs 10% vs 15% in q < 0.05, q < 0.01, q < 0.005, Figure 4A; represented by “I+IV”, Figures 4C and 4E; represented by “I”). No false negatives were identified in both Dups and No Dups at either q < 0.01 or q < 0.005 (Figure 4C and 4E). To determine the downstream impact of having only 96 distinct qRNA-seq labels in single end reads versus 96×96 in paired end reads, we repeated the paired end analysis using only the label of the first sequence in the template. The 96×96 labels had identified an additional 3.6% of duplicates to be non-PCR artifacts (Figure 2C), and this resulted in 4% (55 vs 59%) less Dups false positives at q < 0.05 (Figures 4A and 4B), 10% (35 vs 45%) at q < 0.01 (Figures 4C and 4D), and 10% (42 vs 52%) at q < 0.005 (Figures 4E and 4F).

**Figure 4.**
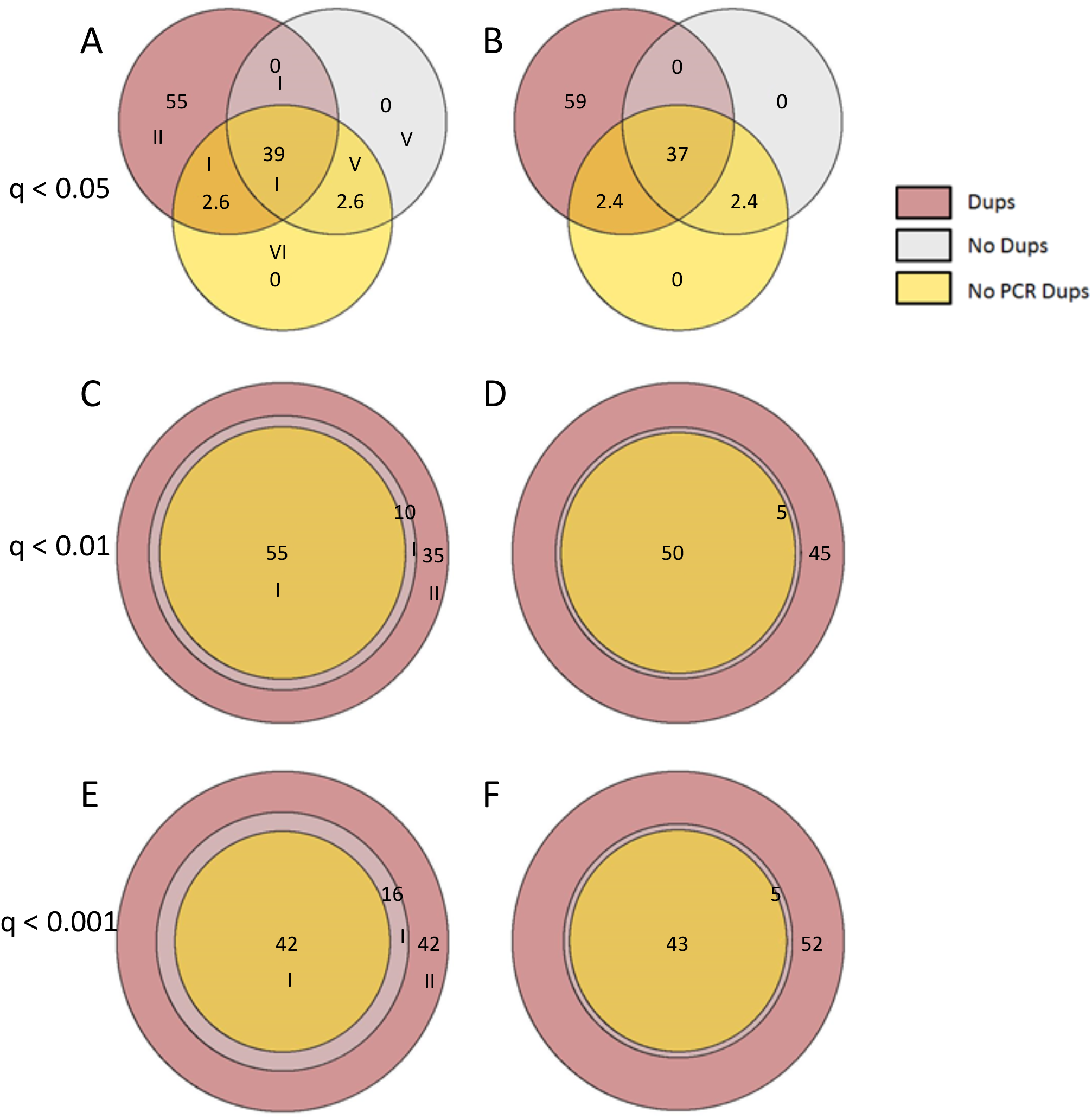
The set of significantly differentially expressed genes at (**A,B**) q < 0.05, (**C,D**) q < 0.01 and (**E,F**) q< 0.005 from the paired end sequences comparison was calculated for 3 alignments: all duplicates retained (Dups), all duplicates removed (No Dups) and only PCR duplicates removed (No PCR Dups). Dups false positives are calculated as sum of regions I+III, No Dups false positives are as sum of regions I+VI, Dups false negatives are as sum of regions V+VII, and No Dups false negatives are as sum of regions IV+VII, respectively. The results were compared by having only 96 distinct qRNA-seq labels in single end reads (**B,D,F**) versus 96x96 in paired end reads (**A,C,E**).

### False findings are affected by transcript length, number of exons and GC content

To investigate the properties of false positives and false negatives, we extracted the significantly differentially expressed genes at q < 0.05 for the single end read comparisons and used the biomaRt R package to obtain their properties, including transcript length, number of exons and percentage of GC content. We found that the false positives of the alignment retaining all duplicates (“Dups”, Figure 5; purple lines) were enriched 1.51-fold for transcripts with length <= 700 nucleotides (Figure 5A) and 1.52-fold for transcripts with 1 exon (Figure 5B), when compared to the true positives set from the alignment retaining only non-PCR duplicates (“True Positive”, Figure 5; pale blue lines). Conversely, the false negatives of this alignment (Figure 5; green lines) were depleted by 1.26-fold of transcripts of short length in the range 200-1400 nucleotides (Figure 5A), and of transcripts with a small number of exons (1.56-fold for 1 exon and progressively smaller depletions as the number of exons increased down to 1.26-fold for 7 exons, Figure 5B). RNA-seq differential expression has been shown to be a function of gene length due to shorter genes having larger variance than longer ones as a result of FPKM normalization [20], which may explain our observations.

**Figure 5.**
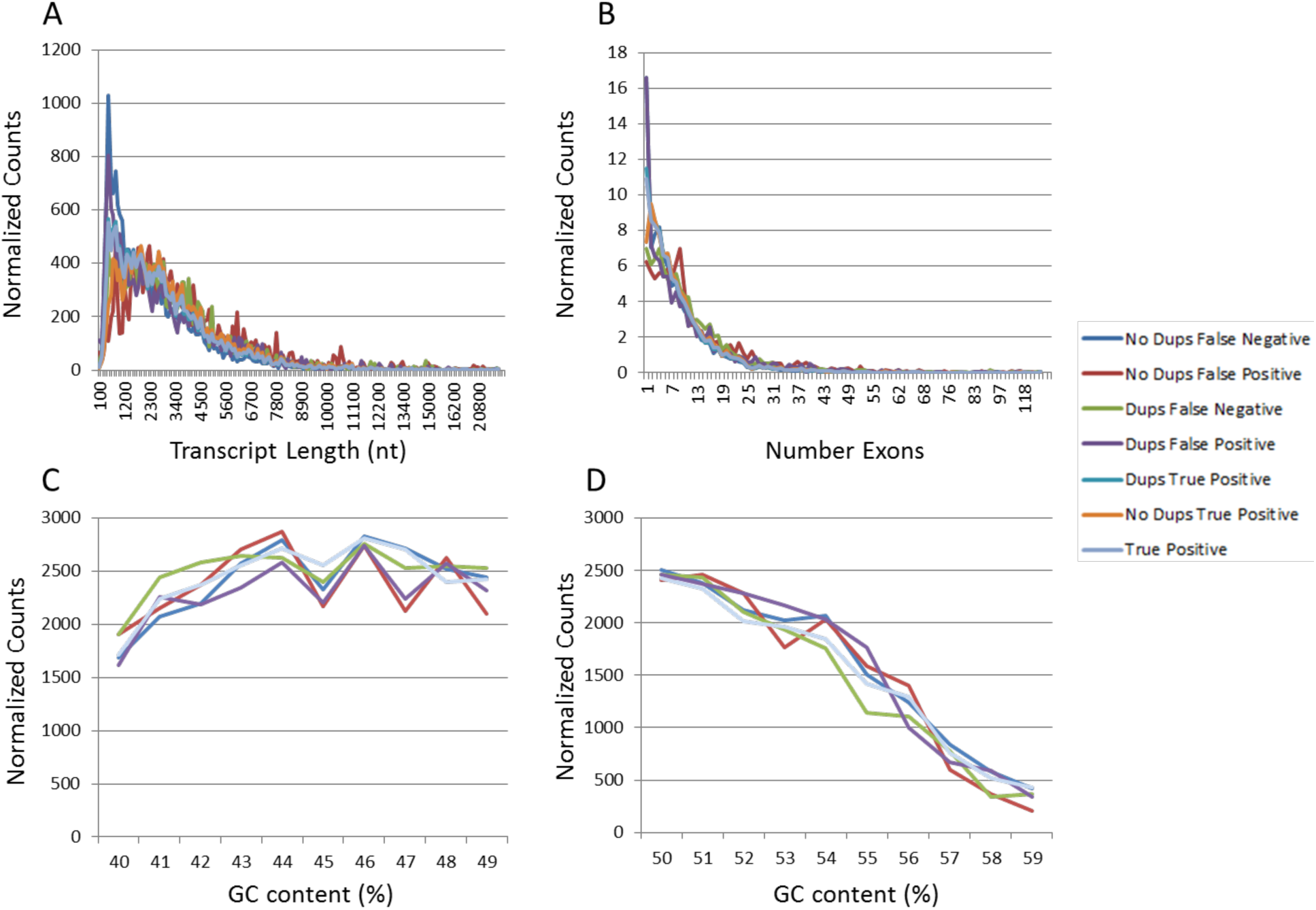
Characterization of the significantly differentially expressed isoforms at q < 0.05 in the true or false, and positive or negative sets of the alignments retaining (Dups) or discarding (No Dups) all duplicates, when taking the corresponding alignments discarding only PCR duplicates (True Positive) as the gold standard. Histograms of A) transcript lengths binned at 100 nucleotides, B) number of exons and C-D) percentage of GC content binned at 1 percent were calculated.

An opposite pattern was observed with the alignment discarding all duplicates (“No Dups”, Figure 5; red and blue lines). In this case, the false positive set (Figure 5; red lines) had a 1.99-fold depletion in the short transcripts with length in the range 200-1400 (Figure 5A), and a 1.75-fold depletion for 1 exon which progressively reduced to 1.37-fold depletion for 7 exons (Figure 5B). The false negative set for this alignment (Figure 5; blue lines) had a 1.64-fold enrichment for transcripts with length <= 700 (Figure 5A) and a 1.52-fold enrichment for transcripts with 1 exon (Figure 5B). This suggests that removing all duplicates, including valid biological ones, is lowering the statistical power of short transcripts increasing their numbers in the false negative set but with the benefit that their number is also decreased in the false positive set. There were also differences in percent of GC content (Figure 5C-D). Since 92% of true positive transcripts had GC content in the range 40-59%, we will restrict our discussion to this range, although there were even bigger differences in the composition of the false negative and false positive isoform sets with respect to the true positives set in GC content outside of this range. The false positive set of the alignments retaining (purple) or discarding (red) all duplicates, and the false negative set of the alignment discarding all duplicates (blue) had less transcripts with GC content in the 40-49% range (5.9%, 3.0% and 1.4% respectively), and more transcripts with GC content in the 50-59% range (4.6%, 0.9% and 4.7% respectively) than the transcripts in the true positive set. The false negative set of the alignment retaining all duplicates (green) had a reverse relation with a 1.9% increase in transcripts with GC content in the 40-49% range and a 3.8% reduction in transcripts with GC content in the 50-59% range (Figure 5C-D).

### Individual qRNA-seq labels did not contribute to the PCR artifacts

To further investigate the potential contributing sources of PCR artifacts, we extended our analysis of qRNA-seq labels. We had previously found that there were many-fold differences in frequencies of individual qRNA-seq labels in the raw fastq sequences that were highly consistent across samples. To determine whether these differences were caused by PCR artifacts, we analyzed the qRNA-seq labels of the PCR and non-PCR duplicates at the nucleotide level. Every qRNA-seq label has 4 A+T and 4 G+C nucleotides, presumably so designed to avoid the long known bias in which PCR templates having GC-rich permutations in the priming site being consistently amplified better than their AT-rich counterparts [18]. Analysis revealed slight differences between the labels in the PCR and non-PCR duplicates alignments in both their positional nucleotide composition and in their dinucleotide frequencies. Specifically, G+C was slightly enriched in the second and third positions of the 8 nucleotide labels in the PCR duplicates with respect to the non-PCR duplicates in each of the 36 single end samples, by 0.71% and 0.45% on average respectively. In addition, of the 16 possible dinucleotides, GT showed the biggest differential enrichment at 0.11% on average, slight enrichment that was present in each of the 36 samples. To determine whether these enrichments were constrained to the labels, we analyzed the transcripts of the true positive, false positive and false negative sets. The start of the transcripts, based on nucleotides 4-10 in order to filter out the first codon, was slightly enriched for G+T in the false positive sets of both the alignments removing or retaining all duplicates when compared to the true positives set (0.60% and 0.36% respectively), indicating that such slight enrichment was not due to PCR artifacts. The dinucleotide GT did not show differences between the transcript sets, and in fact was the third least frequent dinucleotide of the 16 possible dinucleotides in all transcript sets. These findings suggest that the source of the many-fold differences in frequencies of individual qRNA-seq labels in the raw fastq sequences do not originate from PCR artifacts, but rather from steps prior to PCR amplification in the qRNA-seq protocol.

In summary, up to 69% of the single end and 19% of the paired end RNA-seq alignment duplicates originated from valid biological transcripts rather than PCR artifacts. For paired end reads at q < 0.05, the alignment retaining all duplicates resulted in substantial false positives in the sets of significantly differentially expressed genes when considering as the gold standard the corresponding gene set resulting from the alignment having only the PCR duplicates removed, and either removing or retaining all duplicates resulted in a small percentage of false negatives. For single end reads, either removing or retaining all duplicates resulted in a substantial number of false positives and false negatives. Furthermore, the percentage of false negatives and false positives increased as the significance threshold was made progressively more stringent. These data suggest that it is sufficient to remove duplicates from paired end read alignments, but that the qRNA-seq technology greatly improves the sensitivity and specificity of single end RNA-seq downstream analyses.

## Discussion

There are emerging new technologies to mitigate the complications of conventional RNA-seq. The common underlying idea of such technologies is how to avoid PCR amplification. For example, NanoString Technology has eliminated enzymatic reactions and instead employs fluorescence-labeled probes that are uniquely designed for each transcript that can be hybridized and digitally quantified. Oxford Nanopore also eliminated the PCR amplification step and sequenced single cDNAs or RNA molecules directly [15, 16, 17]. It is of great interest how these technologies compete with the qRNA-seq and future investigations are expected to standardize the analytical method to meet the researchers’ specific needs in quantifying transcripts of varying magnitude of abundance. However, technologies employing PCR amplification are still the most commonly used, and here we analyze the impact of PCR artifacts in downstream analyses.

We found that up to 69% of the single end and 19% of the paired end RNA-seq alignment duplicates originated from valid biological transcripts rather than PCR artifacts. For single end reads, either removing or retaining all duplicates resulted in a substantial number of false positives (up to 47.0%) and false negatives (up to 12.1%) in the sets of significantly differentially expressed genes. This gives a warning to all existing gene expression analyses that have utilized single end sequencing methods in short read NGS platforms.

We examined the potential impact of these PCR artifacts by calculating the number of falsely detected- or not detected- significantly differentially expressed genes using ‘real-world’ biomedical samples that comprised of seven phenotypical groups. For single end reads, removing all duplicates resulted on average in 24.8% (15.1-47.0%) false negatives and 4.56% (2.76-6.80%) false positives, and retaining all duplicates resulted on average in 7.35% (3.86-22.0%) false negatives and 8.12% (4.60-12.1%) false positives, at q < 0.05, when considering as the gold standard the corresponding gene set resulting from the alignment having only the PCR duplicates removed. Therefore, without access to, these data confirm the intuitive notion that duplicates should be removed to minimize false positives, and duplicates should be retained to minimize false negatives.

For paired end reads, only the alignment retaining all duplicates resulted in a substantial number of false positives. This indicates that PCR duplicate identification by qRNA-seq technology provided minor benefits for paired end reads. Therefore, we propose that paired end sequencing combined with No Dups analysis approach, in which one will exclude all duplicated reads, can be considered as the most reliable method if not using PCR duplicate identifying technology such as qRNA-seq.

Paired end sequencings, however, is often as twice as expensive as single end analysis. For cost effective gene expression quantification studies using single end NGS platform, qRNA-seq can provide greatly improved sensitivity and specificity in the determination of the significantly differentially expressed genes.

We also found that, for paired end reads, the 96x96 labels identified an additional 3.6% of duplicates to be non-PCR artifacts (i.e. biological replicates) when compared to using only the label of the Read 1 in the template, which resulted in 4% less false positives at q < 0.05 in the alignment retaining all duplicates. This shows that the superior performance of paired end versus single end reads was due mostly to the more stringent definition of read duplication which requires both reads in the template to have identical alignments, rather than the availability of additional labels. This 3.6% difference also shows that the random selection of qRNA-seq labels was biased, since otherwise on average only 1 in 96 (1.04%) additional non-PCR artifacts would have been identified with the addition of a second label. In fact, the distribution of chosen labels was not uniform with enrichment of the most frequent label (“GGCGTATT”) of 1.91 and 1.85-fold, and depletion of the least frequent label (“TAGCTAGC”) of 3.66 and 4.07-fold in the rabbit and mouse samples respectively. This qRNA-seq label selection bias affects the error estimation of the qRNA-seq technology in general. Under unbiased conditions, the resolution of the qRNA-seq technology would be 1/96 (1.04%) for single end reads and 1/(96^*^96) (0. 011%) for paired end reads. However, the qRNA-seq label selection bias measured in this study suggests that the error rate of the qRNA-seq technology is closer to 3.6% for single end reads and 3.6*3.6% or 0.13% for paired end reads.

The false positives of the alignment retaining all duplicates, including PCR artifacts, had 5.9% less transcripts with GC content in the 40-49% range and 4.6% more transcripts with GC content in the 50-59% range. In fact, nucleotide based PCR bias has long been observed, with PCR templates having GC-rich permutations in the priming site found to be consistently amplified better than their AT-rich counterparts. The explanation given for this observation was that since G and C form a triple hydrogen bond, the melting temperatures of the GC-rich permutations of both primers are about 2°C higher, so that at each annealing step a greater proportion of the templates hybridize to their matched primers [18]. More recently, this higher melting property has been shown to result in the depletion of extremely high GC-rich fractions (over 76% or 84%, depending on the heating and cooling rates of the PCR thermocycler used) due to steep thermosprofiles not leaving sufficient time above this critical threshold temperature, causing incomplete denaturation and poor amplification of the GC-rich fraction [19]. We would suggest that GC-bias correction using e.g. EDASeq [21] to be beneficial for differentially expressed gene analysis. Also, any orthogonal method such as qRT-PCR should be included to validate expressions of genes with skewed GC contents or gene length properties.

## Methods

### Animal

All protocols were approved by, and all methods were performed in accordance with the guidelines of, the Institutional Animal Care and Use Committee (IACUC) of Penn State Hershey and Guide of The Association for Assessment and Accreditation of Laboratory Animal Care International (Frederick, MD). For rabbit studies, New Zealand White (NZW) rabbits were maintained in the animal facility of the Pennsylvania State University College of Medicine. Before viral or DNA challenge, rabbits were anaesthetized with a mixture of 40 mg/kg ketamine and 5 mg/kg xylazine. Rabbit backs were shaved and scarified using a scalpel blade, the number of sites depending upon the experiment as described previously [8]. The scarified sites were about 1 cm in diameter and were created by scraping the scalpel blade across the skin to create a “brush burn”-like lesion sufficient to produce a serous fluid with minimal bleeding. Three days following the scarification, the animals were scarified and infected with 5μg of wild type cottontail rabbit papillomavirus (CRPV) DNA or different mutants [9]. For mouse studies, HSD outbred nude (Foxn1^nu^/^nu^) mice (6-8 weeks) were obtained from Harlan Laboratories (ENVIGO). All animals were housed in sterile cages within sterile filter hoods and were fed sterilized food and water in the COM BL2 animal core facility. Mice were sedated i.p. with 0.1ml/10g body weight with ketamine/xylazine mixture (100mg/10mg in10mls ddH_2_O). For tail and muzzle infection, the sites were scarified using a scalpel blade as described previously [8]. One day following the scarification, the animals were scarified and infected with 10μl (1.4 × 10^8^) of the sterilized viral suspension at each site. For vaginal infection, mice were inoculated subcutaneously with 3mg Depo-Provera (Pfizer) in 100 μl PBS three days before the viral infection as described previously. Depo was not administered for anal and oral infections. The vaginal and anal tracts were wounded with Doctors’ Brush Picks coated with Conceptrol (ortho options, over the counter) [10]. Twenty-four hours after wounding, the mice were again anesthetized and challenged with 25μl (3.5 × 10^8^) and 10μl (1.4 × 10^8^) of the sterilized viral suspension at the vaginal and anal tracts respectively. For tongue infection, tongues were withdrawn using a sterile forceps and microneedles were used to wound both the dorsal and ventral surfaces of the tongues. Some bleeding may occur but care was taken to minimize the bleeding. The following day, each animal was again anesthetized. Tongues were again gently abraded and 10μl of sterile virus (1.4 × 10^8^) was applied to the freshly abraded surfaces (10μl each for the dorsal and ventral surfaces) [11]. Animals were placed on their backs during recovery to minimize loss of virus from the infection sites. Monitoring was conducted weekly. The cutaneous lesions and infected mucosal tissues were harvested at different time post infection and stored in Liquid nitrogen.

### RNA sequencing

Total RNA was extracted from both non-infected and infected tissues using mirVana^™^ miRNA Isolation Kit (Thermo Fisher Scientific). RNA integration number (RIN) was measured using BioAnalyzer (Agilent) RNA 6000 Nano Kit to confirm RIN above 7 [12,13]. The cDNA libraries were prepared using the NEXTflex^™^ Illumina qRNA-Seq Library Prep Kit (BioO Scientific) following the manufacturer’s instruction’s. The final product was assessed for its size distribution and concentration using BioAnalyzer High Sensitivity DNA Kit (Agilent). Pooled libraries were diluted (Qiagen) and then denatured using the Illumina protocol. The denatured libraries will be loaded onto a TruSeq Rapid flow cell on an Illumina HiSeq 2500 and run for 64 cycles using a single-read recipe or 100 cycles using a paired-end recipe according to the manufacturer’s instructions. De-multiplexed sequencing reads passing the default purify filtering of Illumina CASAVA pipeline (released version 1.8) were extracted.

### Bioinformatics analysis

The 8 nucleotide qRNA-seq labels were trimmed from the sequences using the FASTX-Toolkit v.0.0.13 (http://hannonlab.cshl.edu/fastx_toolkit). The reads were then aligned to the mouse reference genome (mm10) or the rabbit reference genome (oryCun2) using Tophat v.2.0.13 [14]. The duplicates were annotated using Broad Institute’s Picard MarkDuplicates v.1.102. For single end reads all alignments were extracted and for paired end reads only those with templates having multiple segments in sequencing and where each segment was properly aligned according to the aligner. For single end reads the 8 nucleotide qRNA-seq labels of the aligned reads were recovered from the sequence fastq files based on read name, and for paired end reads both 8 nucleotide qRNA-seq labels, from the right and left sequences of the template, were recovered. Only alignments with valid qRNA-seq labels were kept based on the manufacturer’s list of 96 qRNA-seq labels. For single end reads, PCR duplicates were considered to be the alignments marked by MarkDuplicates as duplicates that had identical chromosome, start, end (start plus read length), cigar string and qRNA-seq label to some other aligned read. For paired end alignments, PCR duplicates had the additional requirement that those fields be identical in both segments of the template and that the template have the same length. In addition, to calculate the impact of the second qRNA-seq label in paired end reads, an alternate set of PCR duplicates were calculated in the same way except that the second qRNA-seq label was not required to match. The RNA transcripts of all reads, non-duplicated reads, and non-PCR duplicated reads were then assembled and their abundances in FPKM values were calculated using Cufflinks v.2.2.1, with a supplied reference annotation (Ensembl gene annotation release 80 for the mouse and the UCSC oryCun2 gtf gene annotation for the rabbit samples) [14]. The mouse and rabbit transcripts were separately merged using Cuffmerge, and mouse groups were compared using Cuffdiff. The significant differentially expressed genes at thresholds of q < 0.05, q < 0.01 and q < 0.005, and isoforms at threshold of q < 0.05 were extracted for each comparison. The biomaRt v.2.30.0 R package was used to obtain isoform annotation from the Ensembl mart and mouse dataset, including transcript length, number of exons and percentage of GC content. A histogram of transcript lengths binned at 100 nucleotides, exons, and percentage of GC content binned at 1 percent were calculated for various sets. The Venn diagrams were calculated using the R packages sets v.1.0-16 and VennDiagram v.1.6.17. Boxplots were generated using the R library sfsmisc v1.1-0.

## Declarations

### Availability of data and materials

The datasets generated during and/or analysed during the current study are not publicly available since the biological finding from these datasets are yet to be published but are available from the corresponding author on reasonable request.

## Footnotes

**Author’s contributions** ACS analyzed the data. ACS and YIK formulated the conception of the study, interpreted the results, and were major contributors in writing the manuscript. EJC, JH, NMC, RMB and GVB performed the sample preparation and NGS analysis. JH provided critical feedback to the manuscript. All authors read and approved the final manuscript.

## Acknowledgements

We thank Neil Christensen, Ph.D. at the Department of Pathology for sharing his RNA-seq samples for this study. We thank Karla Balogh at the Department of Pathology for animal work. We thank Patricia Mantelo at the Department of Pharmacology for assisting with the sample processing. We also thank James Broach, Ph.D. at the Institute for Personalized Medicine and Department of Biochemistry and Molecular Biology and Kent Vrana, Ph.D. at the Department of Pharmacology for mentorship and financial support.

